# Adaptive Protein Evolution in Animals and the Effective Population Size Hypothesis

**DOI:** 10.1101/033043

**Authors:** Nicolas Galtier

**Affiliations:** Institut des Sciences de l’Evolution Université Montpellier – CNRS – IRD – EPHE Montpellier, France

## Abstract

The rate at which genomes adapt to environmental changes and the prevalence of adaptive processes in molecular evolution are two controversial issues in current evolutionary genetics. Previous attempts to quantify the genome-wide rate of adaptation through amino-acid substitution have revealed a surprising diversity of patterns, with some species (*e.g. Drosophila*) experiencing a very high adaptive rate, while other (*e.g*. humans) are dominated by nearly-neutral processes. It has been suggested that this discrepancy reflects between-species differences in effective population size. Published studies, however, were mainly focused on model organisms, and relied on disparate data sets and methodologies, so that an overview of the prevalence of adaptive protein evolution in nature is currently lacking. Here we extend existing estimators of the amino-acid adaptive rate by explicitly modelling the effect of favourable mutations on non-synonymous polymorphism patterns, and we apply these methods to a newly-built, homogeneous data set of 44 non-model animal species pairs. Data analysis uncovers a major contribution of adaptive evolution to the amino-acid substitution process across all major metazoan phyla – with the notable exception of humans and primates. The proportion of adaptive amino-acid substitution is found to be positively correlated to species effective population size. This relationship, however, appears to be primarily driven by a decreased rate of nearly-neutral amino-acid substitution due to more efficient purifying selection in large populations. Our results reveal that adaptive processes dominate the evolution of proteins in most animal species, but do not corroborate the hypothesis that adaptive substitutions accumulate at a faster rate in large populations. Implications regarding the factors influencing the rate of adaptive evolution and positive selection detection in humans *v*s. other organisms are discussed.

**Author summary:** The rate at which species adapt to environmental changes is a controversial topic. The theory predicts that adaptation is easier in large than in small populations, and the genomic studies of model organisms have revealed a much higher adaptive rate in large population-sized flies than in small population-sized humans and apes. Here we build and analyse a large data set of protein-coding sequences made of thousands of genes in 44 pairs of species from various groups of animals including insects, molluscs, annelids, echinoderms, reptiles, birds, and mammals. Extending and improving existing data analysis methods, we show that adaptation is a major process in protein evolution across all phyla of animals: the proportion of amino-acid substitutions that occurred adaptively is above 50% in a majority of species, and reaches up to 90%. Our analysis does not confirm that population size, here approached through species genetic diversity and ecological traits, does influence the rate of adaptive molecular evolution, but points to human and apes as a special case, compared to other animals, in terms of adaptive genomic processes.

## Introduction

Characterizing and quantifying adaptation at the molecular level is one of the major goals of evolutionary genomics. Recent research in this area has yielded a number of remarkable examples [1,4]. However, in spite of the increased size of molecular data sets and intensive methodological developments, we still lack a quantitative assessment of the relative impact of adaptive *vs*. nearly-neutral evolution at the genomic scale – an issue hotly debated ever since the origins of molecular evolutionary studies [5–7].

Particular attention has been paid to the adaptive evolution of proteins trough amino-acid substitution, following the seminal work of McDonald and Kreitman [8]. Analysing the *Adh* gene in *Drosophila*, these authors compared the ratio of the number of non-synonymous (potentially selected) to synonymous (supposedly neutral) substitutions between species, *d_N_/d_S_*, to the ratio of the number of non-synonymous to synonymous polymorphisms within species, *p_N_/p_S_*. They showed that the former ratio was higher than the latter, which was interpreted as resulting from adaptive evolution. This is because the rare, positively selected non-synonymous mutations contribute negligibly to the polymorphism and to *p_N_/p_S_*, but have a much higher fixation probability than neutral mutations, thus accumulating detectably over time as species diverge, and inflating *d_N_/d_S_*. In absence of adaptive evolution, *d_N_/d_S_* is expected to be equal to *p_N_/p_S_* (neutral model), or lower than *p_N_/p_S_* (nearly-neutral model [9]).

The idea that the excess of *d_N_/d_S_* over *p_N_/p_S_* is a signature of adaptive protein evolution was turned into methods for estimating the proportion of amino-acid substitutions that result from positive selection, a quantity called α [10–15]. These methods differ in the way information is combined across genes, and in the way they cope with the two main confounding factors that have been identified, i.e., (i) slightly deleterious mutations, which tend to inflate *p_N_/p_S_* while not affecting *d_N_/d_S_*[16], and (ii) variation in time of effective population size (*N_e_*), which tends to decouple *d_N_/d_S_* from *p_N_/p_S_* irrespective of adaptive processes [17].

In their most sophisticated version, McDonald-Kreitman-based methods model the deleterious effects of non-synonymous mutations by assuming some (typically Gamma or log-normal) distribution of selection coefficients, recent changes in *N_e_* being accounted for either parametrically or non-parametrically. The model is fitted to polymorphism data, typically depicted by the synonymous and non-synonymous site-frequency spectra (SFS) [18,19]. Slightly deleterious mutations, which confound the McDonald-Kreitman signal, are expected to segregate at lower frequency, on average, than neutral mutations, thus distorting the non-synonymous SFS, compared to the synonymous one. SFS-based estimates of population genetic parameters are used to predict the expected divergence process under near-neutrality, and specifically the expected *d_N_/d_S_* ratio, which, compared to the observed one, leads to the estimation of α [15].

These methods have been applied to various species in which large numbers of protein-coding sequences were available from several individuals and a closely related outgroup. Interestingly, the resulting estimates of α varied considerably among species. For instance, the estimated α was close to zero in apes [14,20], giant Galapagos tortoise [21], yeast [22] and nine plant species [23], but above 0.5 in *Drosophila* [11], mouse [24], rabbit [25], sea squirt [26], sunflower [23] and enterobacteria [27]. According to these results, positive selection would be the prevalent mode of protein evolution in some species, but a minor one in others, including humans, in which the vast majority of amino-acid substitutions are predicted to be nearly neutral and fixed through genetic drift. Such a contrasted pattern among species is in itself intriguing, and deserves an explanation.

The main hypothesis that has been considered so far regarding across-species variation in the molecular adaptive rate invokes a population size effect. Several lines of evidence suggest that α is higher in large-*N_e_* than in small-*N_e_* species. The hypothesis first arose from the human *vs*.

*Drosophila* comparison [15, 28], and was somewhat corroborated by the report of a low α in typical small-*N_e_* species [21] and a high α in typical large-*N_e_* ones [26]. In sunflower [29] and house mouse [30], the comparison of closely related species revealed a greater α in the most highly polymorphic species, which is consistent with the *N_e_* hypothesis. Finally, a meta-analysis of 13 eukaryotic species revealed a significant – albeit not so strong – positive relationship between *N_e_* and the rate of adaptive substitution [31]. Not all the empirical evidence, however, is consistent with the *N_e_* hypothesis [23, 32], the most obvious counter-examples being provided by the yeast and maize proteomes, which, despite presumably very large *N_e_*, are apparently dominated by non-adaptive processes.

Theoretically, there are mainly two reasons why large*-N_e_* species are expected to experience a higher adaptive rate than small-*N_e_* ones [28]. First, the population rate of appearance of new beneficial mutations is proportional to *N_e_*, i.e., higher in large populations. Secondly, once they have appeared, beneficial mutations have a greater probability of fixation in large populations than in small populations, in which the random effect of genetic drift is stronger. These arguments have been debated [6,7], however, and two recent simulation studies using Fisher’s geometric formalism recovered only a weak relationship between *N_e_* and the adaptive rate [33, 34].

The determinants of the variation in α among species and the relevance of the effective population size hypothesis therefore remain contentious, in large part because we still lack a broad overview of the prevalence of adaptive processes in nature. First, the sample of species for which we have an estimate of α, which are mostly model organisms, is limited and disparate. Secondly, the set of analysed genes varied much between studies. For instance, in humans, more than 10,000 protein-coding sequences have been analysed [14], whereas the above mentioned plant study was based on a few dozens to a few hundreds of loci [23]. Finally, estimates of *a* and ω_a_ have been obtained using distinct methods and models across studies, each making specific assumptions about the evolutionary processes at work.

Here, we analyse patterns of coding sequence polymorphism and divergence in the high-expressed genomic fraction of 44 taxonomically and ecologically diverse metazoan species characterized by a substantial variation in *N_e_* [35]. A battery of models for the distribution of fitness effects of mutations (DFE) are introduced and assessed in the maximum likelihood framework, with particular emphasis on the modelling of beneficial mutations. The confounding effects of sample size, polymorphism orientation errors, gene expression level and GC-content are considered and controlled for. The analysis revealed a pervasive impact of adaptive processes on the evolution of coding sequences in the eight sampled animal phyla, but did not detect any influence of *N_e_* on the rate of adaptive molecular evolution.

## Results

Forty four metazoan species pairs were analysed (S1 Table). Each pair consisted of a focal species, in which at least four diploid individuals were available, and a closely-related outgroup species. The data set was based on two recent transcriptome-based population genomic reports [35, 36]. Five species were newly sequenced in this study (one individual each). The sample covered eight metazoan phyla: 18 pairs of vertebrates (of which twelve mammals), eleven arthropods (of which eight insects), five molluscs, four echinoderms, two tunicates, one annelid, one nematode, one nemertian and one cnidarian. Focal species sample size varied from four to eleven individuals, and was seven or more in 26 species. Ten of the twelve mammalian species had a sample size of four [36]. The number of analysed coding sequences varied between 295 (Black and white ruffed lemur *Varecia variegata variegata*) and 7951 (small skipper *Thymelicus sylvestris*), the median being 1900. The total number of single-nucleotide polymorphisms (SNPs) varied between 315 (*Varecia variegata variegata*) and 62,553 (Essex skipper *Thymelicus lineola*), the median being 3241. The number of synonymous SNPs and the number non-synonymous SNPs were both above 100 in all the analysed species. The per site synonymous divergence between focal and outgroup species, *d_S_*, varied between 0.7% (in penguins *Aptenodytes patagonicus* vs. *A. forsteri*) and 36% (in nematodes *Caenorhabditis brenneri* vs. *Caenorhabditis* sp.16).

### DFE model comparison

In all pairs of species, a population genetic model was fitted to the synonymous and non-synonymous polymorphism (*i.e*., SFS) and divergence data. Model parameters included population mutation rate θ, demography *sensu lato* **r**, neutral divergence *T*, and the distribution of fitness effects of non-synonymous mutations (DFE). The 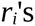 parameters – one per allele frequency class – multiply both the synonymous and the non-synonymous SFS; they are intended to capture any departure from the mutation/selection/drift equilibrium (see Methods and reference [19]). Six distinct parametrizations of the DFE were used. These included the classical Neutral[11] and Gamma [19] models, which only consider negative selection coefficients, and four newly introduced models explicitly accounting for beneficial mutations (see Methods). DFE models were compared using the Akaike Information Criterion (AIC) and likelihood ratio tests.

Among the 44 species pairs, the Neutral model obtained the highest AIC score in four cases, and the Gamma model in 13 cases. In the remaining 27 cases, *i.e*., 60% of the data sets, a model accounting for slightly advantageous mutations best fitted the data. The GammaExpo model, which assumes a Gamma distribution of negative effects and an exponential distribution of positive effects, obtained the highest AIC score in 18 data sets. The ScaledBeta model, which uses a Beta-shaped distribution of mild negative and positive effects, obtained the highest AIC score in 8 data sets. The DisplacedGamma model [37] essentially converged towards Gamma, the displacement parameter always taking very low values. The BesselK model [38] sometimes yielded a likelihood similar to (but not above) GammaExpo, and sometimes performed badly – perhaps due to optimization problems. These two models were not considered any longer in this study. When we performed likelihood ratio tests between the nested Neutral, Gamma and GammaExpo models, we found that Gamma significantly rejected Neutral in 40 species, and GammaExpo significantly rejected Gamma in 19 species out of 44.

Consistent with published methods [15], the above-described approach implicitly assumes the existence of strongly advantageous non-synonymous mutations that contribute to divergence but negligibly affect polymorphism patterns (parameter A in equation (7), see Methods). Since the GammaExpo and ScaledBeta models account for the beneficial effect of non-synonymous mutations, we implemented a distinct version, the [-A] version, in which no additional, strongly adaptive class of mutation was assumed. This was done for the 26 data sets in which either GammaExpo or ScaledBeta was the model best supported by AIC. In 14 of these, the [-A] version was not significantly rejected by the classical one according to likelihood-ratio test. In these cases the new DFE models seem to appropriately account for both polymorphism patterns and divergence rates. In the remaining twelve data sets, the fit was significantly worse in the [-A] than in the classical version. Estimates of α were strongly correlated across species between the two analyses (*n*=44, r^2^=0.79, *p*-val<10^−10^).

### Adaptive substitution rate

The estimated population genetic parameters were used to predict the expected rate of non-adaptive non-synonymous substitutions relative to neutral, ω_na_, deduce the estimated rate of adaptive ones, ω_a_= *d_N_/d_S_* − ω_na_, and the proportion of adaptive non-synonymous substitutions, α=ω_a_/(*d_N_/d_S_*) (see Methods). This was achieved using four distinct DFE models. Fig. 1 plots the Neutral, GammaExpo and ScaledBeta estimates of α as a function of the Gamma estimate. The Neutral model provided estimates generally lower than, and loosely correlated with, the other three models. This confirms that accounting for the effect of weakly deleterious non-synonymous mutations makes a significant difference, as previously demonstrated [14–16]. The GammaExpo and ScaledBeta estimates of α, in contrast, were strongly correlated with the Gamma estimate, suggesting that the inclusion of positive effects in the assumed DFE does not impact much the estimation of the adaptive rate, even when a better fit to the data is achieved (Fig. 1, closed circles). Below we focus on the estimate of α based on the GammaExpo model, but very similar results would be obtained with either the Gamma of ScaledBeta estimates. Maximum likelihood confidence intervals for α are provided in S1 Table.

**Figure 1.**
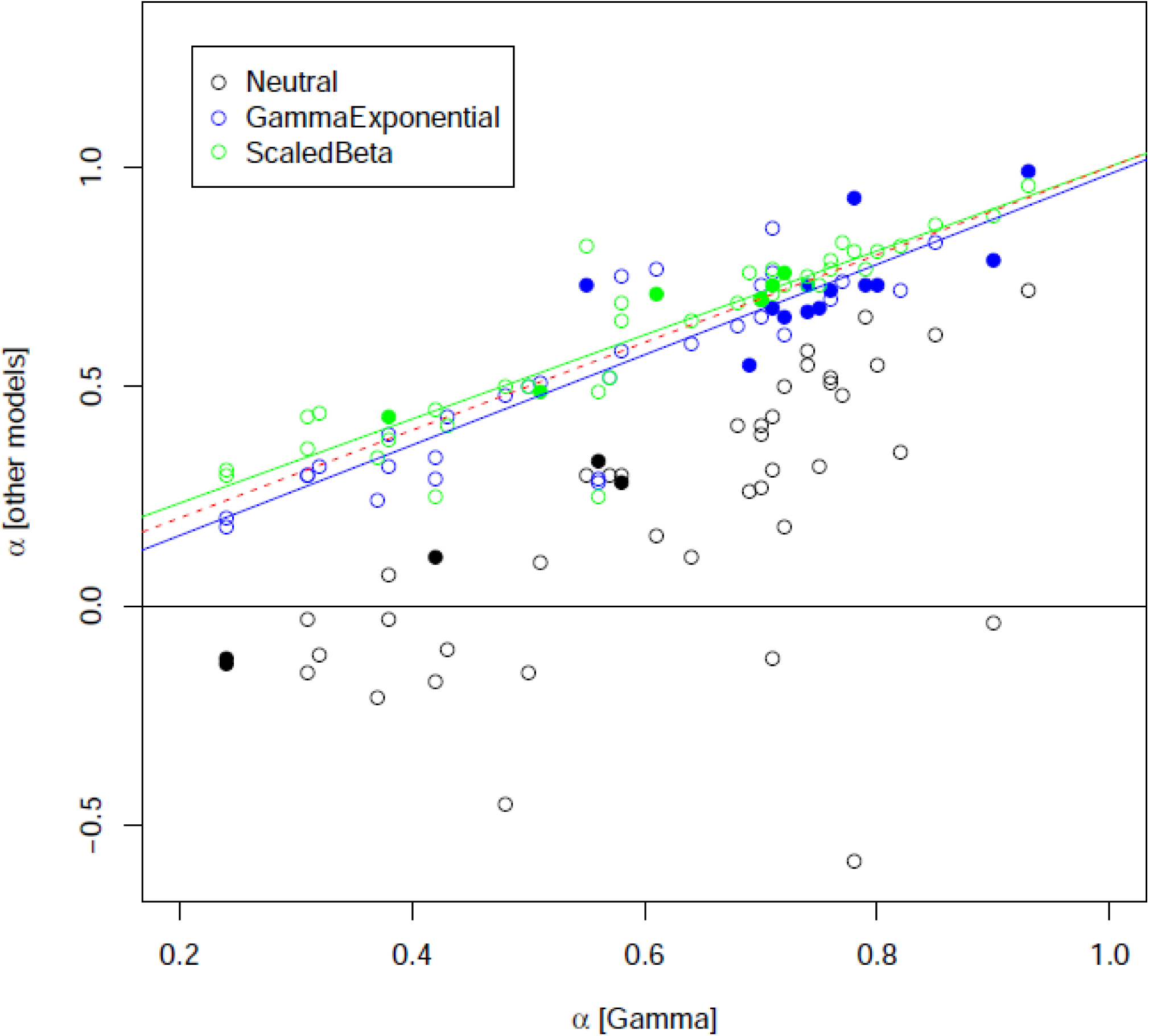
Effect of the underlying DFE model on estimators of α. Each dot is for a species pair. X-axis: estimated α under the Gamma model. Y-axis: estimated α under the Neutral (black), GammaExpo (blue) and ScaledBeta models (red). Dotted red line: (y=x) line. Closed circles indicate species pairs for which the considered model obtained the highest AIC.

The distribution of α among animal species was bimodal, with a peak around 0.3 and another one around 0.7, whereas the distribution of ω_a_ was unimodal (S1 Figure). No significant difference in α was detected between vertebrates (18 species) and invertebrates (26 species, Fig. 2, top left). Within invertebrates, no difference was observed between arthropods (eleven species), molluscs (five species) and echinoderms (four species). Within vertebrates, however, a surprising significant difference was detected between mammals (twelve species, of which ten primates) and sauropsids (three bird and two turtle species), α being lower than in the average metazoan in the former, but higher in the latter (Fig. 2, bottom left). The mammalian pattern is mainly driven by primates: the two non-primates species pairs yielded relatively high estimates of α (Iberian hare *Lepus granatensis:* α=0.77; common vole *Microtus arvalis:* α=0.60). A more or less similar pattern was found regarding the taxonomic distribution of ω_a_ (Fig. 2, right).

**Figure 2.**
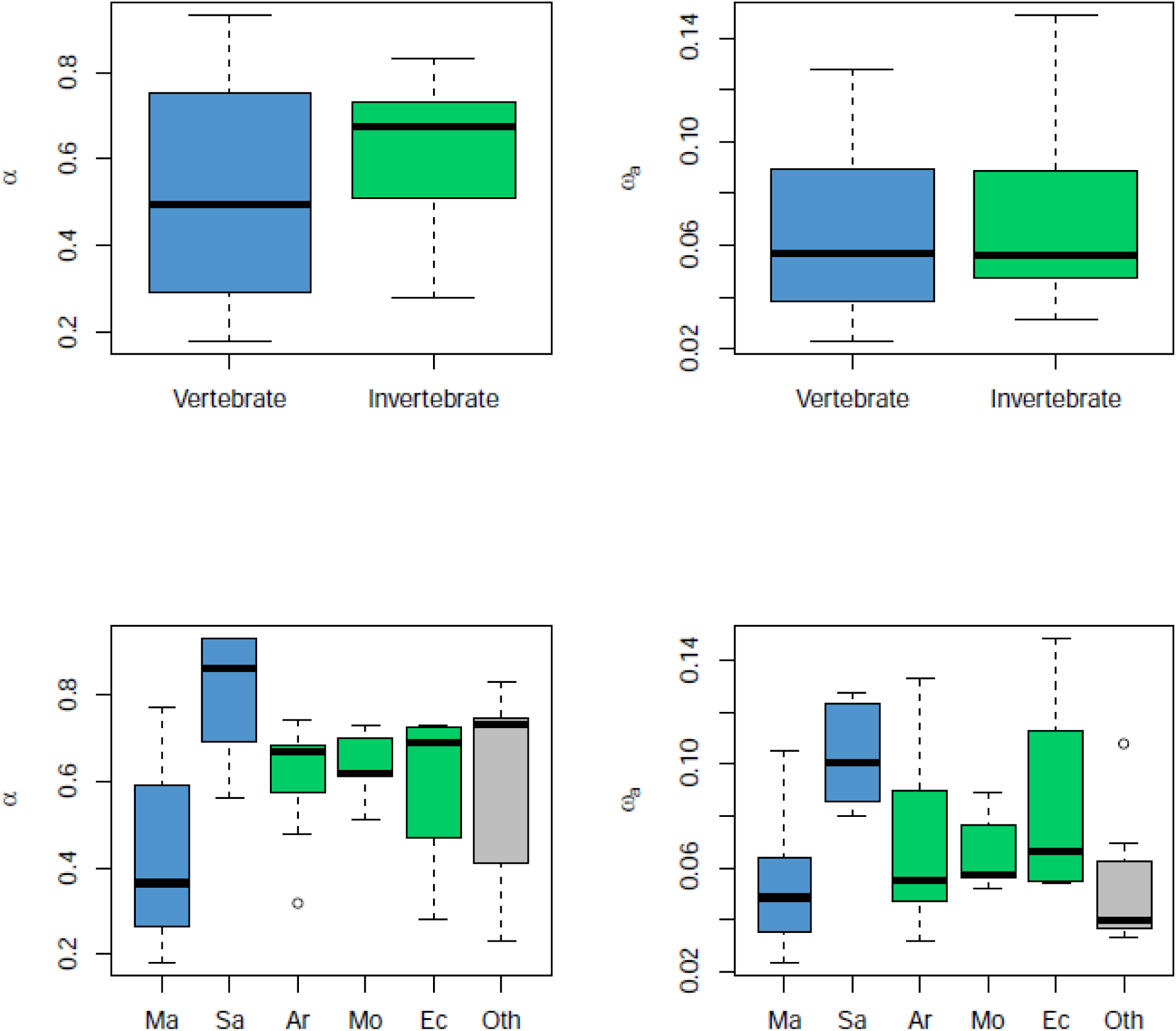
Distributions of α and ω_a_ split by taxa. Ma: mammals; Sa: sauropsids; Ar: arthropods; Mo: molluscs; Ec: echinoderms; Ot: others

We correlated *d_N_/d_S_*, ω_na_, ω_a_, and α to species genetic diversity, π_S_, here taken as a predictor of *N_e_*, the effective population size. A significant negative relationship between ω_na_ and log-transformed π_S_ was obtained (Fig. 3a), which is indicative of a higher efficiency of purifying selection against slightly deleterious mutations in large-*N_e_* species [35]. The relationship between *d_N_/d_S_* and π_S_ was also a significantly negative one (Fig. 3b), suggesting that the effect of *N_e_* on the efficiency of purifying selection also affects the rate of non-synonymous substitution. Interestingly, the slopes of the ω_na_ vs. log(π_S_) and *d_N_/d_S_* vs. log(π_S_) relationships were quite similar, as we illustrated by reporting in Fig. 3b the regression line of Fig. 3a (dotted line). According to the McDonald-Kreitman rationale, the parallel lines indicate that the adaptive component of the non-synonymous rate, ω_a_= *d_N_/d_S_* − ω_na_, is unrelated to *N_e_*, as shown by Fig. 3c. The proportion of adaptive amino-acid substitutions, finally, was significantly and positively related to log-transformed π_S_ (Fig. 3d), consistent with the existing literature. However, Fig. 3a-c suggests that the variation of α among species is primarily driven by the fixation rate of slightly deleterious changes. α tends to be lower in small*-N_e_* than in large*-N_e_* species because non-adaptive substitutions accumulate at a faster rate in the former, thus decreasing the proportion of adaptive ones. The correlation coefficients and *p*-values corresponding to the four regression analyses shown in Fig. 3 are provided in Table 1 (analysis A).

**Figure 3.**
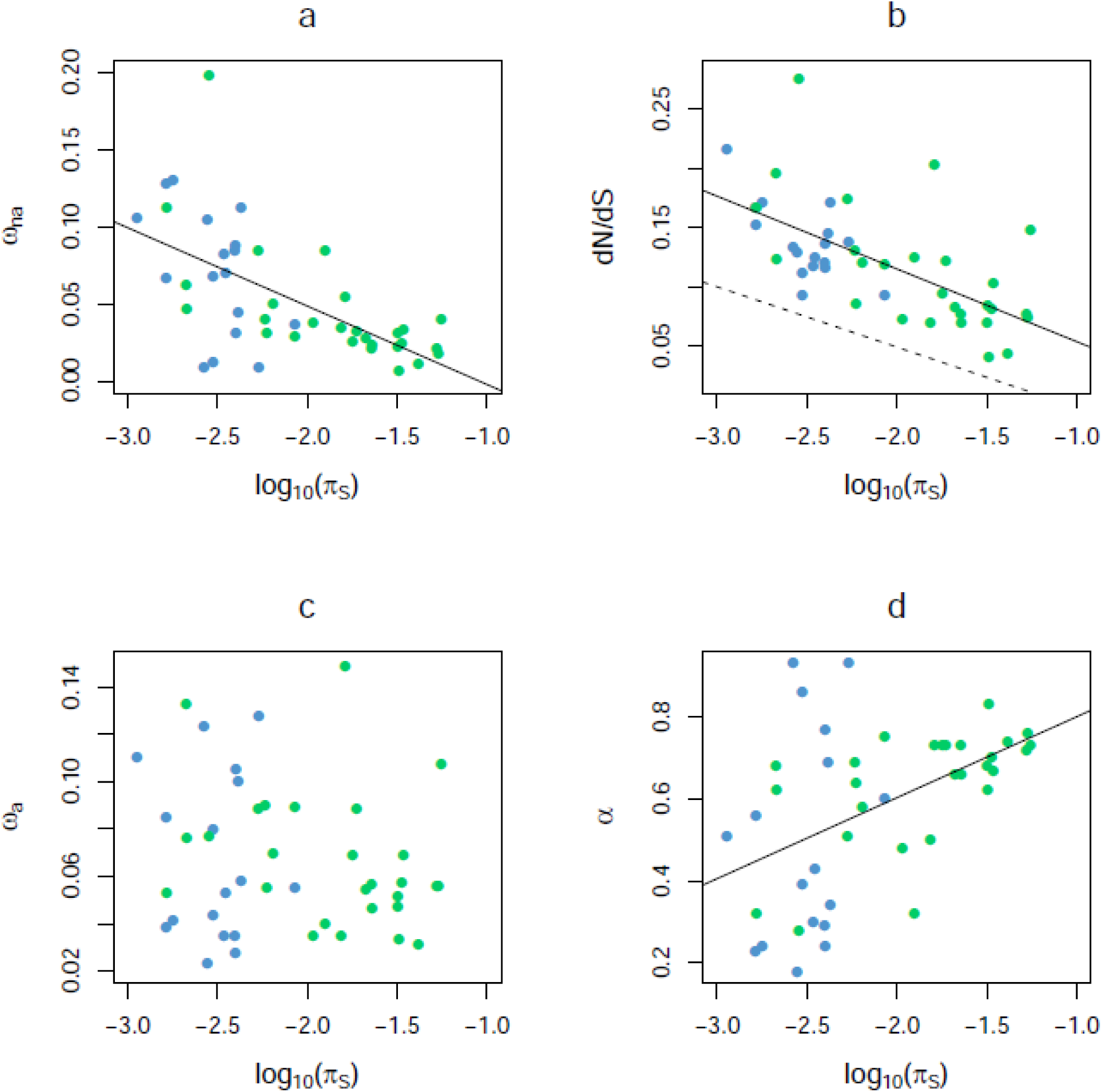
Effect of species neutral genetic diversity on protein divergence parameters. Blue: vertebrates; green: invertebrates.

In Fig. 3a, 3c and 3d the X-axis, π_S_, is not independent of the Y-axis – estimators of α, ω_a_ and ω_na_ depend on the synonymous diversity. We therefore re-conducted the same analysis using other predictors of *N_e_*, namely species maximal longevity, and size of propagule – propagule being defined as the stage that leaves the mother. These two life-history traits were identified as strong negative correlates of *N_e_* in animals [35]. We obtained results very similar to those shown in Fig. 3, *i.e*., a significant correlation with these two traits for *d_N_/d_S_* and ω_na_, but not for ω_a_, α being in this case only marginally significant (S2 Figure, S3 Figure). The proportion of adaptive non-synonymous substitutions tends to be higher in short-lived, small propagule-sized species, but this is due in the first place to a faster accumulation of non-adaptive changes in long-lived, large propagule-sized species. None of the other life-history traits considered in ref. [35] (body mass, fecundity, size of geographic range) did provide additional explanation to the across-species variation in α and ω_a_.

**Table 1.** Genetic diversity vs. protein divergence: control analyses

| analysis | sp.<sup>a</sup> | ind.<sup>b</sup> | $\alpha$ vs. $\pi_s$<sup>c</sup> | $d_N/d_S$ vs. $\pi_s$ | $\omega_{na}$ vs. $\pi_s$ | $\omega_a$ vs. $\pi_s$ |
| --- | --- | --- | --- | --- | --- | --- |
| A: main | 44 | 7.2 | 0.48 ; $<10^{-3}$ | -0.64 ; $<10^{-5}$ | -0.62 ; $<10^{-5}$ | -0.17 ; NS |
| B: >6 individuals | 26 | 9.2 | 0.20 ; NS | -0.70 ; $<10^{-4}$ | -0.48 ; 0.012 | -0.60 ; $<10^{-2}$ |
| C: subsampled 4 | 40 | 4 | 0.49 ; $<10^{-2}$ | -0.57 ; $<10^{-3}$ | -0.61 ; $<10^{-4}$ | -0.10 ; NS |
| D: folded SFS | 26 | 9.2 | 0.11 ; NS | -0.70 ; $10^{-4}$ | -0.49 ; 0.012 | -0.61 ; $<10^{-2}$ |
| E: $d_S$ control | 21 | 7.3 | 0.76 ; $<10^{-4}$ | -0.63 ; $<10^{-2}$ | -0.80 ; $<10^{-5}$ | 0.17 ; NS |
| F: GC-poor | 18 | 8.8 | 0.46 ; NS | -0.50 ; 0.03 | -0.65 ; $<10^{-2}$ | -0.23 ; NS |
| G: GC-medium | 21 | 9.0 | 0.32 ; NS | -0.65 ; $<10^{-2}$ | -0.69 ; $<10^{-3}$ | -0.49 ; 0.024 |
| H: GC-rich | 21 | 9.0 | 0.32 ; NS | -0.65 ; $<10^{-2}$ | -0.55 ; $<0.011$ | -0.51 ; 0.018 |
| I: low-expressed | 23 | 9.0 | 0.17 ; NS | -0.62 ; $<10^{-2}$ | -0.48 ; $<0.021$ | -0.49 ; 0.018 |
| J: high-expressed | 25 | 9.1 | 0.18 ; NS | -0.69 ; $<10^{-3}$ | -0.50 ; 0.012 | -0.56 ; $<10^{-2}$ |
| K: whole body | 22 | 8.4 | 0.57 ; $<10^{-2}$ | -0.69 ; $<10^{-3}$ | -0.79 ; $<10^{-4}$ | -0.38 ; NS |
| L: no mirror | 36 | 7.6 | 0.44 ; $<10^{-2}$ | -0.63 ; $<10^{-4}$ | -0.59 ; $<10^{-4}$ | -0.25 ; NS |

<sup>a</sup> number of analysed species<sup>b</sup> mean number of individuals per species<sup>c</sup> [correlation coefficient ; $p$ -value]; NS: not significant

### Effect of sample size

Our dataset is heterogeneous across species in terms of number of analyzed individuals. We investigated the influence of sample size on our results in two ways. We first re-conducted the above analyses using only the 25 species for which at least eight individuals were available. Similar to the full dataset analysis, we found a significantly negative relationship between *d_N_/d_S_* and log-transformed π_S_, and between ω_na_ and log-transformed π_S_. No significant correlation, however, was detected between α and π_S_ with the large sample-size dataset, and ω_a_ was negatively related to π_S_ (Table 1, analysis B) – an intriguing result that contradicts the *N_e_* hypothesis. It should be noted that taxon sampling is more evenly balanced amongst animal phyla in this 25 species data set, which includes only two mammalian species, than in the complete data set, which includes twelve mammals. In a second analysis, we analysed the whole set of species but used sub-samples of exactly four individuals in each species – only the 40 species for which the number of non-synonymous SNPs was above 100 were retained. The results were again strongly consistent with the main analysis, suggesting that the heterogeneity of sample size is not a major issue in this analysis (Table 1, analysis C).

### SNP mis-orientation

In the above analyses we used so-called unfolded SFS, in which the derived and ancestral alleles are defined based on the allelic state of the outgroup. Orientation errors can deeply affect the folded SFS in that mis-oriented low-frequency mutations are mistaken for high-frequency ones. This problem, however, is alleviated in the current approach by the “demographic” 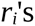 parameters, which are expected to capture any departure from the expected SFS as soon as it is shared by synonymous and non-synonymous sites. Still mis-orientation could be problematic if it did not affect synonymous and non-synonymous sites to the same extent. To control for this potential bias we built folded SFS, in which SNPs in frequency *k* and 2*n-k* are merged in a single bin, *n* being the diploid sample size. Only the 25 species in which sample size was seven individuals or more were considered here. Estimates of α were significantly correlated between the two analyses (*n*=26, r^2^=0.28, *p*-val<10^−2^), but there was a couple of outliers in which the folded and unfolded SFS yielded quite different estimates of α – e.g., 0.78 (folded) *vs*. 0.32 (unfolded) in the yellow gorgonian *Eunicella cavolinii*. Regarding the relationship to π_S_, however, the results with folded SFS were essentially similar to those obtained with this set of species and unfolded SFS (Table 1, analysis D). We also restricted the analysis to a subset of 21 species in which *d_S_* was between 0.05 and 0.25 substitution per synonymous site, thus avoiding biases due to a too high, or a too low, level of divergence. Again, this control yielded results qualitatively similar to the main analysis (Table 1, analysis E).

### GC-content

For each species pair, we split the data set in three bins of equal size according to coding sequence third codon position GC-content (GC3) and re-estimated α separately for GC-poor, GC-median and GC-rich genes. Within each category of genes, the relationships between α, *d_N_/d_S_*, ω_a_, ω_na_ and π_S_ shown in Fig. 3 were recovered (Table 1). Considering only the species in which sample size was seven of more and above 100 non-synonymous SNPs were available, we found a slight, positive effect of GC3 on α (Fig. 4). This might reflect the effect on the efficacy of positive selection of local recombination rate, which alleviates Hill-Robertson effects by breaking genetic linkage – GC3 is associated to local recombination rate in a number of animal species [39,40]. Alternatively, a higher α in high-GC regions might reflect the confounding effect of GC-biased gene conversion, a recombination-associated molecular drive that mimics the effect of directional selection [41].

**Figure 4.**
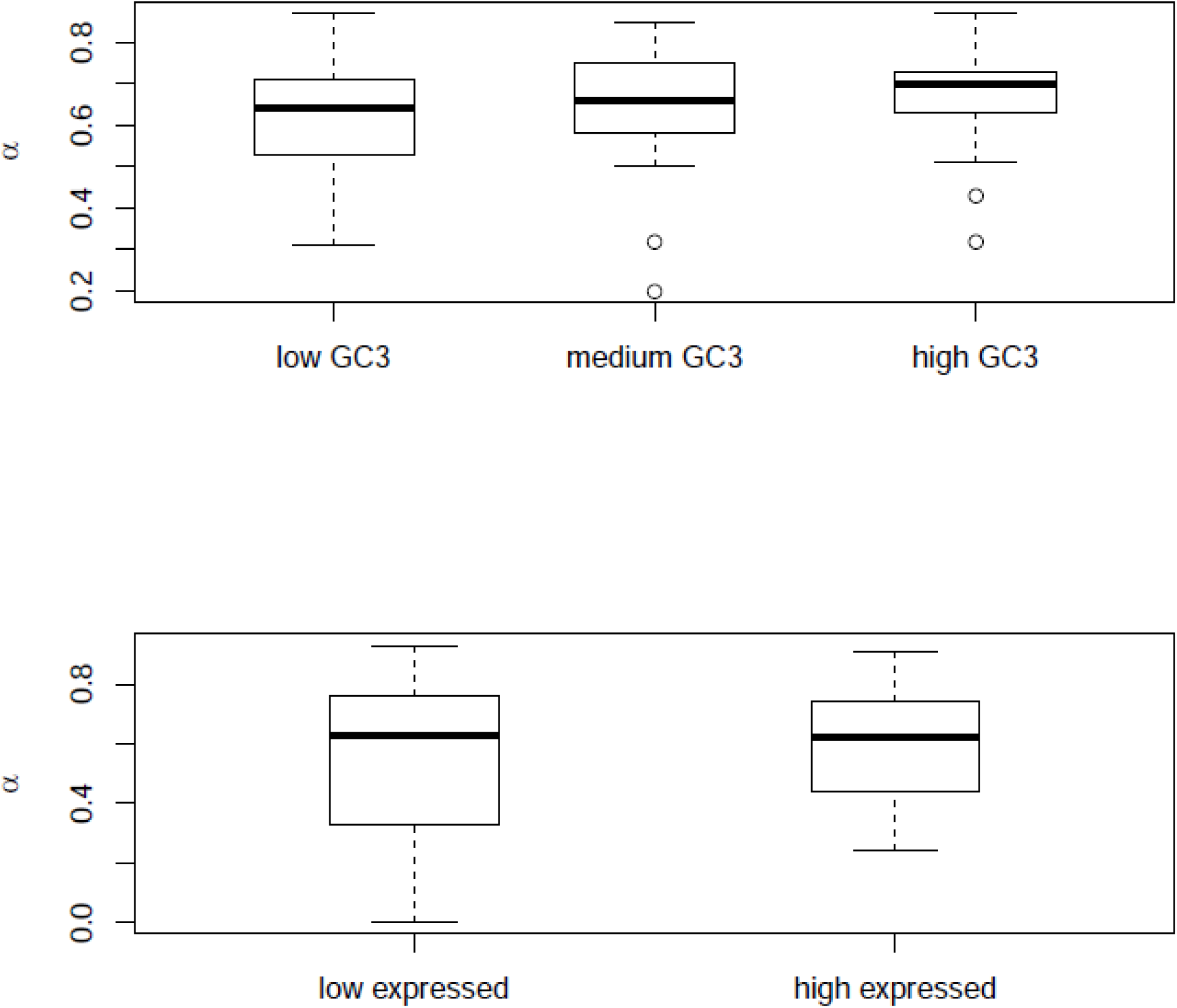
Effect of gene GC-content and expression level on estimates of α. Only the 26 species for which at least seven individuals have been sampled are considered here.

### Gene expression level

For each species pair, we split the data set in two bins according to gene expression level and reestimated a separately for low-expressed and high-expressed genes. In this case bins were unequal in size: the low-expressed bin included 90% of the genes, and the high-expressed bin 10% of the genes. This is because there is a strong positive correlation between ORF length, genotyping success and expression level in this transcriptome-based data set – high-expressed genes are more efficiently assembled and genotyped – so that using bins of equal size would result in a strong discrepancy in terms of number of SNPs. Consistent with the existing literature [42], we detected a significant effect of expression level on the ratio of non-synonymous to synonymous changes: the across-species average π_N_/π_S_ of high-expressed genes (0.084) was lower than that (0.096) of low-expressed genes (*p*-val<0.001, *n*=44, Student’s t-test for paired samples). However, no difference in α was detected between the two categories of gene expression (Fig. 4), which, considered separately, yielded results similar to those of Fig. 3 (Table 1).

### Tissue specificity

The sets of analysed genes varied among species, particularly due to the fact that not the same tissues were used in distinct species for RNA extraction – e.g., blood in reptiles, liver in mammals, intestine in urchins. This might affect the comparison if the prevalence of positive selection was higher in genes expressed in specific tissues. To control for this effect, we analysed the subset of 22 species in which whole-body RNA extraction had been performed. This includes insects, molluscs, annelids, tunicates, nematodes, cnidarians, some echinoderms and some crustaceans. Vertebrates were excluded from the sub-sample, but the range of π_S_ was unaffected. The analysis provided results highly similar to the main one (Table 1, analysis K).

### “Mirror” species

Our data set included eight pairs of “mirror” species, in which species 1 served as outgroup for species 2, and reciprocally (S1 Table). First, when we randomly selected a single species per mirror species pairs, reducing the data set to 36 species, results were essentially unchanged (Table 1, analysis L). Mirror species, on the other hand, are useful because they offer biological replicates: the two analyses (species 1 outgroup to species 2 *vs*. species 2 outgroup to species 1) concern the same history of divergence, and therefore estimate the same biological quantities.

To get a hint about the robustness of estimates of α (and ω_a_), we compared their values between mirror species. In four mirror pairs out of eight the difference in α was below 0.1 between the two mirror analyses, but in three mirror pairs the difference reached values above 0.15 – and up to 0.26 in the *Ciona intestinalis A* / *Ciona intestinalis B* pair. In neither of the eight mirror pairs was the α estimate of species 1 within the confidence interval of α in species 2, and reciprocally. This indicates that the ML confidence intervals (S1 Table), which implicitly trust the assumptions of the model, are unrealistically narrow. Interestingly, in six mirror pairs out of eight, the higher estimate of α was obtained in the more polymorphic of the two mirror species (S4 Figure).

### Characterizing the DFE

Besides α and ω_a_, our model-fitting procedure also provides estimates of parameters characterizing the DFE. When the Gamma model was used, the shape parameter of the negative Gamma distribution of selection coefficients (call it β) varied greatly among species - from 0.009 to its maximum allowed value 100.0 – and took a median value of 0.23. When the GammaExponential model was used, the median β was higher and equal to 0.46. With Gamma distributed negative selection coefficients the theory predicts a log-linear relationship of slope 1- β between π_N_ and π_S_ [43,44]. The linear regression of log(π_N_) on log(π_S_) across species yielded an estimated slope of 0.62 and an estimated β of 0.38, in reasonable agreement with species-specific, SFS-based estimates. These results indicate that the data examined in this study are broadly consistent with a negative Gamma distribution of shape in the 0.2 – 0.4 range for selection coefficients applying to non-synonymous mutations, even though single species data can be too noisy to provide reliable estimates of the DFE.

## Discussion

The comparative approach offers a unique opportunity to relate species traits to population genomics, and particularly investigate the effect of selection and drift on molecular evolutionary processes [35, 45]. In this study, we extended existing McDonald-Kreitman-based approaches to estimate the rate of adaptive amino-acid substitution from coding sequence polymorphism and divergence analysis. We applied the new methods to the high-expressed genomic fraction of 44 metazoan species pairs, thus offering for the first time a general picture of the distribution of the adaptive rate across taxa in this group.

### Enhanced McDonald-Kreitman modelling

Various population genomic models involving distinct shapes for the DFE were implemented in the maximum likelihood framework and assessed through simulations (S1 Text). Including a continuous distribution of positive effects in the model significantly improved the fit in a majority of data sets. In roughly half of these, this addition was sufficient to correctly fit both the polymorphism and divergence data. In the other half, *d_N_*/*d_S_* was still in excess compared to model predictions, thus requiring an extra class of arbitrarily high selection coefficients, similarly to classical methods [15]. It should be noted that the proportion of adaptive non-synonymous mutations and the average strength of adaptive effects could not be efficiently disentangled – the two parameters were strongly, negatively correlated. Similar results were reported in a previous attempt to incorporate adaptive effects in the DFE [46] – in this case, a single, discrete category of positive selection coefficients was assumed. Among the newly introduced DFE models, GammaExpo and ScaledBeta were the ones best fitting the data. The two DFE models that were suggested from theoretical studies based on Fisher’s geometric formalism did not perform well in this analysis. Importantly, the new approaches did not make a strong difference with respect to estimates of α and ω_a_, compared to existing ones.

In this study, departure of the synonymous SFS from the neutral, constant-*N_e_* expectation was accounted for thanks to the “anonymous” 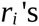 parameters [19]. This is biologically less meaningful but statistically more flexible than explicit models of population size changes [47]. The 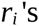 parameters are intended to capture the effects of a variety of potential confounding factors, such as demographic changes, population structure, strong selection at linked loci, genotyping error or SNP misorientation – provided these factors affect synonymous and non-synonymous SNPs to the same extent (S1 Text). This methodological choice was motivated by a recent simulation study [48], which showed that linked selection can distort the synonymous and non-synonymous SFS in a way that confounds explicit models of demographic changes.

### Pervasive adaptive protein evolution in animals

The analysis of 44 species paired revealed a prominent effect of adaptation on amino-acid sequence evolution in animals. Our estimate of α, the proportion of adaptive amino-acid substitutions, was above 0.5 in a majority of the analysed species and taxa – with the notable exception of primates (Fig. 2). Observing a high α therefore seems to be the rule, and a low α the exception, making humans and apes a peculiar, intriguing case. The report of prevalent adaptive protein evolution in animals contrasts with the plant situation, in which eight out of nine analysed species pairs showed little evidence for positive selection [23]. Identifying the environmental and genomic drivers of protein sequence adaptation in animals and understanding the causes of the difference with plants would be two exciting goals to be pursued.

An exciting goal of current population genomics is to uncover the targets of positive selection. Our results suggest that the challenge is even stronger in humans and apes than in most animal species. The confounding effect of neutral and slightly deleterious mutations, which are prevalent in primates, is likely to affect not only *d_N_*/*d_S_*-based studies, but also analyses of local SNP-density [49] and of Fst-outlying loci [50]. The focus of this study is on point mutations in coding sequences, but the results are expected to be general to a wider range of mutations, including structural and regulatory changes, unless the DFE associated to these mutation types differ substantially from that of amino-acid sequence changes [51].

### No effect of N_e_ on the adaptive rate?

A significant, positive effect of various predictors of *N_e_* on α was detected, in broad agreement with the existing literature. This relationship, however, seems to be primarily explained by deleterious effects. Being more strongly affected by genetic drift, small-*N_e_* species accumulate slightly deleterious non-synonymous substitutions at a higher rate than large-*N_e_* ones, in agreement with the nearly neutral theory of molecular evolution ([52,53]). This mechanically decreases the proportion of adaptive non-synonymous substitutions. No positive relationship was found, however, between ω_a_, the (relative to neutral) rate of adaptive amino-acid substitution, and *N_e_*. This result, which was robust to controls for DFE modelling, sample size, divergence time, SNP orientation errors, GC-content, and gene expression level, does not corroborate a recent report of a weak but significant effect of *N_e_* on ω_a_ across 13 species from four eukaryotic groups [31], but confirms earlier suggestions that the long-term adaptive rate does not always respond to changes in *N_e_* [32].

Our results are consistent with theoretical predictions recently obtained by simulations under Fisher’s geometric model [34], in which α was only weakly positively related to *N_e_*, and ω_a_ independent from *N_e_*. A plausible reason for these perhaps counter-intuitive results is that proteins in small-*N_e_* species tend to be less adapted (*i.e*., further away from their fitness optimum), and therefore more prone to adaptation, than in large-*N_e_* species. The accumulation of slightly deleterious amino-acid substitutions in small-*N_e_* species would open the opportunity for compensatory, adaptive ones – at least under the hypothesis of a single-peaked fitness landscape [34], and assuming that mutation does not limit adaptation [54]. A positive relationship between ω_a_ and *N_e_* would be expected if the DFE was independent of *N_e_*, *i.e*., if the adaptive mutation rate was similar in small and large populations. Our study, if confirmed, would suggest that this hypothesis does not hold, at least in animals: adaptive mutations could be more common in small-*N_e_* species, and this would compensate for their lower fixation probability [55,56].

The lack of a strong positive relationship between ω_a_ and *N_e_* does not contradict the prediction that adaptive walks are more efficient, in terms of rate of fitness increase, in large than in small populations [57]. This is because McDonald-Kreitman-related methods estimate the number of adaptive steps, not the associated fitness effect. To our best knowledge, no analytical result is available regarding the relationship between the number of steps in adaptive walks and *N_e_*. Simulations under Fisher’s geometric model and a fluctuating environment, however, seem to indicate that the length of adaptive walks is essentially independent of *N_e_*, at least for the range of parameters considered in references [33] and [34]. In these simulations large populations adapt more efficiently than small populations by making larger steps (on average), not more steps.

Not detecting a positive relationship between ω_a_ and *N_e_* does not demonstrate the absence of such a relationship. The McDonald-Kreitman method is known to be sensitive to various confounding effects, and our data could be just too noisy (see below methodological discussion). However, what our analysis reveals is a significantly negative relationship between *d_N_*/*d_S_* and *N_e_* (Fig. 3b). This indicates that *N_e_* primarily affects the rate of amino-acid sequence evolution through its effect on slightly deleterious mutations. The effect of *N_e_* on the adaptive rate, if any, must be of second order – or *d_N_*/*d_S_* would correlate positively with *N_e_*.

In this study, like in previous ones [23, 29, 30], the synonymous diversity, π_S_, was used as a proxy for *N_e_*. This can be problematic for several reasons. First, π_S_ is influenced not only by *N_e_* but also by the mutation rate. This can in principle be addressed by incorporating direct or indirect estimates of the per generation mutation rate in the analysis [31]. Unfortunately, data on generation time and divergence time are lacking for many of the species analysed here. The strong relationships between π_S_ and ω_na_ and between π_S_ and *d_N_*/*d_S_*, however, suggest that *N_e_* is the major determinant of π_S_, since ω_na_ and are *d_N_*/*d_S_* independent of the mutation rate (and see [35]). Secondly, synonymous sites are not always neutrally evolving and might be affected by selection on codon usage. The robustness of our results to variation in GC-content, which is typically correlated to codon usage bias, does not suggest that this is a major issue with this analysis. Thirdly, some of the species analysed here are highly polymorphic (*e.g. C. brenneri*), and the standard formula for π_S_ does not hold in such extreme cases [58]. Figure 3, however, does not reveal any particular effect of the most highly polymorphic species on the main results of this analysis.

### What determines the between-species variation of ω_a_?

The evolutionary genomic literature in animals has long been dominated by studies in humans (and apes) and *Drosophila*. Quite plausibly, the patterns that distinguish these two groups have often been associated to variation in *N_e_*. Our analysis of a taxonomically extended data set suggests that this is not obviously the right explanation as far as the adaptive rate is concerned, re-opening the question of the determination of the variation of the adaptive rate across species. We report a wide range of estimated α (minimum: 0.18; maximum: 0.93) and ω_a_ (minimum: 0.023; maximum: 0.15) among animals, without obvious connection to species biology or taxonomy. Several hypotheses can be considered.

First, it might well be that distinct species do not face the same rate of environmental change. Unfortunately, measuring or predicting the rate at which abiotic and biotic environmental fluctuations affect the fitness landscape of distinct species appears essentially impossible, so that this hypothesis appears difficult to test empirically. Secondly, a part of the variance of α and ω_a_ might be explained by the fact that different sets and numbers of genes have been analysed among species – even though the data set analysed here is arguably more homogeneous than in previous meta-analyses. Gene expression levels vary substantially among metazoan phyla, and, depending on the species, RNA was extracted from either the whole body or specific tissues [35], so that distinct sets of genes were captured. The prevalence of adaptive evolution is known to vary across gene sets and functions, with a enrichment among adaptive loci of genes associated to the immune response [49, 59]. Results were unchanged, however, when we restricted the analysis to species in which whole-body RNA extraction was performed. Clarifying the influence of gene function on estimates of α and ω_a_ would be a promising continuation of this work; this should preferably be achieved based on whole proteomes in model organisms, in which functional annotations are available.

Finally, the dispersion of α and ω_a_ estimates among species could be in part due to methodological issues. First, the data are somewhat noisy, as suggested by the wide variation across species in estimated shape of the DFE, and by the higher variance in estimated α among low-diversity species, compared to high-diversity ones (Fig. 3d). Secondly, it might be that not all the sources of biological variation are appropriately modelled in the McDonald-Kreitman framework, as previously suggested [60, 61]. This includes local adaptation, balancing selection and other sources of protected selected polymorphisms, which if common might push π*_N_*/π*_S_* above its nearly-neutral expectation. Illustrative of shortcomings of the model, the likelihood-based confidence intervals we obtained are implausibly narrow, compared to the variation resulting from slight modifications of the data analysis strategy – for instance, switching from unfolded to folded SFS substantially impacted the estimate of α in some species. One particularly critical assumption is the time-constancy of *N_e_* during the whole period of divergence. Our model accounts from recent demographic changes through the *ri* ‘s parameters, but events older than the coalescence time of the sampled individuals, or having appeared in the outgroup lineage, might have influenced the *d_N_*/*d_S_* ratio while being undetectable from current polymorphism patterns. A former bottleneck, for example, might have inflated *d_N_*/*d_S_* through the accumulation of slightly deleterious non-synonymous substitutions in a way indistinguishable from truly adaptive evolution. Our simulations, however, suggest that the approach is sufficiently robust to detect large differences in α and ω_a_ between species in case of fluctuating *N_e_* (S1 Text).

Polymorphism data only provide information on current, or very recent, strength of purifying selection, whereas *d_N_*/*d_S_* is influenced by the long-term average. Our analysis of “mirror” species, in which distinctive estimates of α were obtained depending on which species was taken as the focal one, highlights limitations of the McDonald-Kreitman approach when based on polymorphism data from a single species. The advent of genome-wide polymorphism data in multiple, closely related species should help disentangling the effects of demography *vs*. positive selection on the evolution of amino-acid sequences.

## Methods

### Data sets

The data set was built based on two recent population genomic studies of, respectively, 16 mammalian [36] and 76 metazoan [35] species. In these two contributions, two to eleven individuals per species were sampled from distinct populations and subjected to massive mRNA sequencing. From this sample we identified 35 focal species for which (i) at least four individuals had been sequenced, (ii) a closely-related outgroup was available, and (iii) at least 100 synonymous SNPs and 100 non-synonymous SNPs had been called. We added nine outgroup species (one individual each), of which four were already published [21,62,63] and five were newly sequenced here, increasing the total sample size to 44 species pairs (species list and SRA IDs in S1Table). Not all these species pairs are fully independent from each other: when four individuals or more were available in each of two closely species, two species pairs were defined, hereafter referred to as “mirror species pairs”, each species serving as outgroup to the other one. A restricted data set of 36 species pairs was created by randomly keeping just one mirror species pair out of two.

The newly-sampled individuals were subjected to RNA extraction, library preparation and illumina sequencing as previously described [64]. In each of the analysed species, sequence reads were assembled to predicted cDNAs and open reading frames (ORFs), then single nucleotide polymorphisms (SNPs) and diploid genotypes were called the same way as in [35], according to the methods described in previous papers of ours [26,62,65]. For each species pair, orthologous coding sequences between the focal and outgroup species were predicted by reciprocal best-BLAST hit, aligned using MACSE [66] and cleaned as previously described [63].

For each species pair, the numbers of fixed synonymous, *D_S_*, and fixed non-synonymous, *D_N_*, differences were computed. In each focal species, the synonymous, **P_S_**, and the non-synonymous, **P_N_**, unfolded SFS were built by computing the observed frequency of derived alleles, and summing counts across genes. An unfolded SFS is a vector of 2*n*-1 entries corresponding to the counts of SNPs at which the absolute frequency of the derived allele is 1, 2, …, 2*n*-1, respectively, where *n* is the sample number of diploid individuals. SNP orientation was deduced from the outgroup state. SNPs for which the outgroup was polymorphic or distinct from any of the focal species alleles were discarded, as well as tri-allelic and tetra-allelic SNPs. For each species pair, the data set consisted of 4*n* data points: *D_N_*, *D_S_*, and the 2*n*-1 entries of each of **P_N_** and **P_S_**. Capital letters are used here to denote counts, lower case *d_N_*, *d_S_*, **P_N_** and **P_S_** being used for rates – *i.e*., counts divided by site numbers.

Not every position of every individual was genotyped, so that sample sized varied among SNPs, complicating the calculation of the SFS. To cope with missing data we arbitrarily defined *n*, the minimal required number of genotyped diploid individuals, and applied a hypergeometric projection of the observed SFS into a subsample of size *n*, SNPs sampled in less than *n* individuals being discarded [67]. We used *n*=4 when total sample size was four or five individuals, *n*=5 when total sample size was six individuals, *n*=6 when total sample size was seven or eight individuals, and *n*=7 when total sample size was above eight individuals.

### Population genetic model

We used a modified version of Eyre-Walker’s approach [15,19] to fit a population genetic model to the data. Consider a focal species of population size *N* that diverged *t* generations ago from its sister species (outgroup). Mutations occur at rate µ per site per generation and drift applies according to the Wright-Fisher model under panmixy. Synonymous mutations are assumed to be neutral, whereas non-synonymous mutations experience protein-level codominant selection with constant in time selection coefficient *s* against heterozygotes, *s* being drawn from distribution φ(*s*) – the DFE. The expected number of synonymous and non-synonymous SNPs at derived allele frequency *i*, 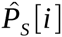 and 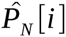, can be expressed as:

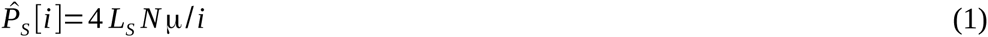

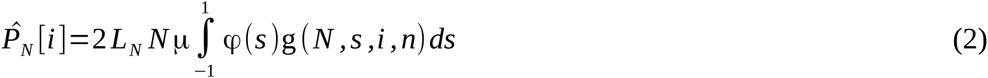

where *L_S_* and *L_N_* are the sampled numbers of synonymous and non-synonymous sites, respectively.

In equation (2), g(*N*, *s*, *i*, *n*) is the probability that a mutation of selection coefficient *s* segregates at observed frequency *i* in a sample of size 2*n*. The latter can be expressed by integrating on allele frequency, *x*:

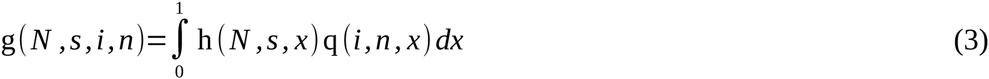

where h(*N*, *s*, *x*) is the relative sojourn time at frequency *x* of a mutation of selection coefficient *s*, and q(*i*, *n*, *x*) is the binomial probability that an allele at population frequency *x* is observed at frequency *i* in a sample of size 2*n*.

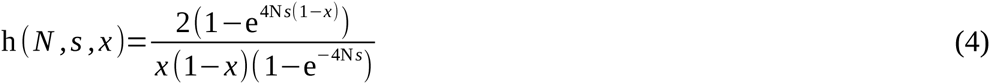

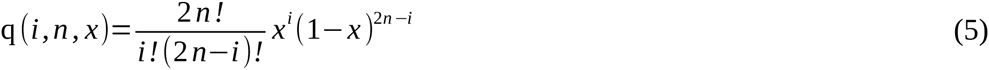

### Departure from the Wright-Fisher assumption

The shape of observed SFS’s typically differ from the expectations of equations (1) and (2). There are many potential reasons for this, including population sub-structure, variation in time of *N*, and interference between linked selected mutations. Following [19], we used a generalized version of equations (1) and (2):

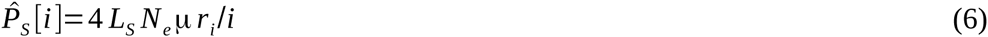

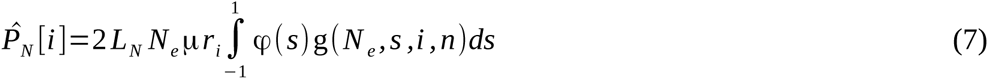

in which *r*_1_=1 and {*r*_2_, *r*_3_, …, *r*_2n-1}_ are intended to capture any effect, typically demographic, affecting both the synonymous and non-synonymous SFS. Equating all the 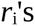 coefficients to unity would model the case of a constant-size Wright-Fisher population. The population size parameter in equations (6) and (7) is now called “effective” and denoted by *N_e_*. It represents the size of a Wright-Fisher population that would confer the same amount of distortion between non-synonymous and synonymous SFS as observed in current data. Accordingly, *N* should be replaced with *N_e_* in equations (4) and (5) as well.

### Divergence model

The expected number of synonymous and non-synonymous substitutions, 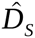 and 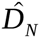, were expressed as:

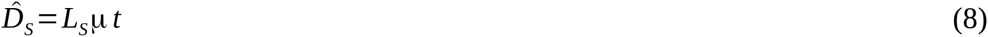

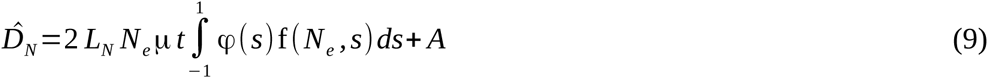

The first term in equation (9) corresponds to the expected number of non-synonymous substitutions given µ, *t*, *N_e_*, *s*, and φ .f(*N_e_*, *s*)=2*s*/(1-exp(-4 *N_e_ s*)) is the fixation probability of a mutation of selection coefficient *s* in a population of size *N_e_*. The additional parameter *A* accounts for the excess of non-synonymous substitutions due to strong positive selection. Introducing parameter *A* here is equivalent to adding to the DFE a class of mutations of arbitrarily high selection coefficient that contribute to divergence but negligibly affect polymorphism patterns.

Not all model parameters are identifiable since *N_e_*, µ, *s* and *t*, which appear as products in the above equations, capture just three degrees of freedom. We used the following re-parametrization:

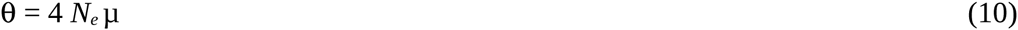

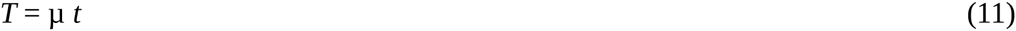

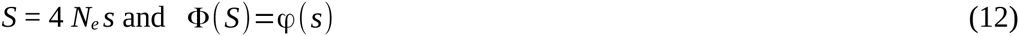

### DFE models

Several functions were used to model the distribution of fitness effects of non-synonymous mutations. The Neutral model considers a discrete DFE, with a class of neutral mutations and a class of strongly deleterious mutations, which do not contribute to polymorphism or divergence [8,11]. The Gamma model, in contrast, assumes the existence of weakly deleterious non-synonymous mutations, modelled as a continuous, negative Gamma distribution [19].

Besides these classical DFE, we investigated models assuming a continuous distribution of positive effects. The GammaExpo model builds upon Gamma by adding a proportion of weakly advantageous mutations, assumed to be exponentially distributed. The DisplacedGamma model assumes a displaced negative Gamma distribution [37], and the BesselK model implements the DFE obtained by Lourenco et al (their equation 8) [38]. These two DFE derive from two distinct versions of Fisher’s geometric model. Finally, the ScaledBeta model is similar to the Neutral model in including a class of strongly deleterious mutations, but assumes a Beta-shaped distribution of weak-effect mutations:

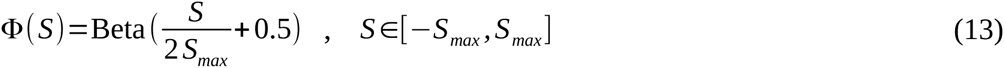

Here *S*_max_ was arbitrarily set to 25. The Beta distribution has two parameters and can take many different shapes – monotonously increasing, monotonously decreasing, flat, unimodal, U-like.

### Maximum likelihood estimation and the adaptive amino-acid substitution rate

Parameters θ, Φ, **r**, T and *A* were fit to the data using the maximum likelihood method. The likelihood was calculated assuming a Poisson distribution of polymorphism and substitution counts:

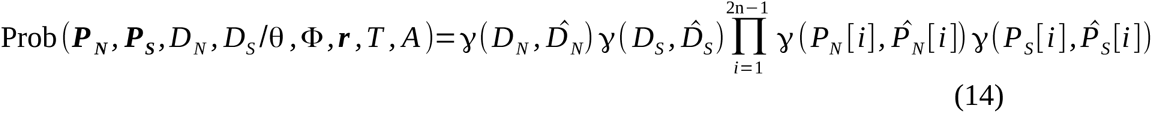

where 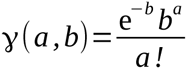

Based on these parameter estimates we calculated 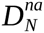, the expected number of non-adaptive (*i.e*., nearly-neutral) non-synonymous substitutions. Non-synonymous substitutions were said to be non-adaptive when corresponding to a mutation of population selection coefficient below threshold *S*_adv_:

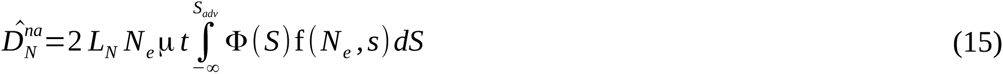

*S*_adv_ was here arbitrarily set to *S*_adv_=5. For the Neutral and Gamma DFE models, this is equivalent to the existing methods [11, 15].

Comparing this expectation to the actual number of non-synonymous substitutions and dividing by the observed number of synonymous substitutions, we calculated ω_a_, the rate of non-adaptive amino-acid substitution relative to neutral divergence, ω_a_, the rate of adaptive amino-acid substitution relative to neutral divergence, and α, the proportion of adaptive non-synonymous substitutions:

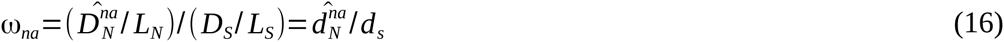

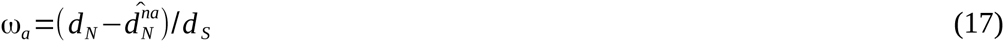

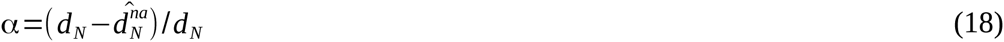

Alternatively, for DFE models explicitly accounting for beneficial mutations, we used a version in which *A* was set to zero, so that no additional class of strong selection coefficient was assumed. This version, which was called [-A], has one less degree of freedom than the classical. Comparing the two versions is a way to test whether the divergence pattern is appropriately predicted by polymorphism data and the model, or if an extra parameter is needed. Models were compared using likelihood-ratio tests (LRT) when nested, and Akaike’s Information Criterion (AIC) otherwise. Integrals in equations (3), (7), (9) and (15) were calculated numerically. The likelihood was maximized using the Newton-Raphson method. Confidence intervals around estimates of α were defined as values of α for which the log-likelihood was within two units of its maximum. A C++ program implementing these methods was written based on the Bio++ libraries [68]. Source code and Linux executable files are available upon request. The reliability of α and ω_a_ estimates obtained under the Gamma, GammaExponential and ScaledBeta models were assessed from simulations (S1 Text). Linear regressions were performed in R using Pearson’s parametric method.

## Acknowledgements

The author is grateful to Thomas Bataillon, Sylvain Glémin for their help with the theoretical part, Philip Messer for help with his simulation program, the Montpellier Bioinformatics & Biodiversity platform for computing facilities, Dmitri Petrov and four anonymous reviewers for thoughtful comments.

## Supporting Information Caption

**S1 Table.** Species pairs, polymorphism and divergence data, α and ω_a_ estimates, confidence intervals. Pn, Ps, Dn, Ds refer to the total number of non-synonymous and synonymous SNPs and substitutions, respectively. Lpn, Lps, Ldn, Lds refer to the total number of non-synonymous and synonymous sites available in the within-species and between-species alignments, respectively. α_down and α_up refer to the confidence interval boundaries for α (and similarly for ω_a_). Estimates were obtained under the GammaExpo model.

**S1 Figure.**
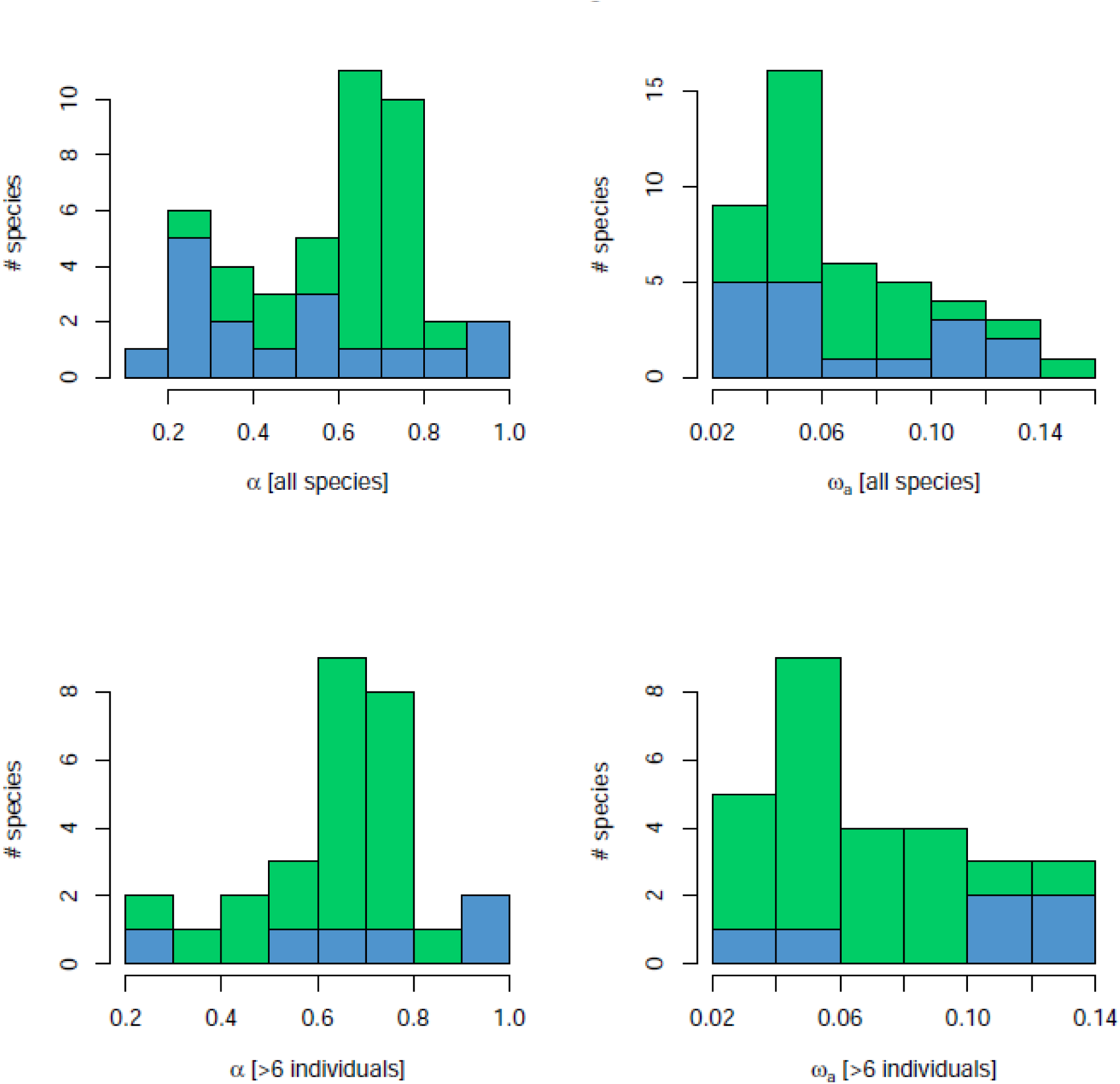
Distribution of α and ω_a_ estimates across 44 animal species pairs. Top: all species; Bottom: 26 species for which at least seven individuals have been sampled.

**S2 Figure.**
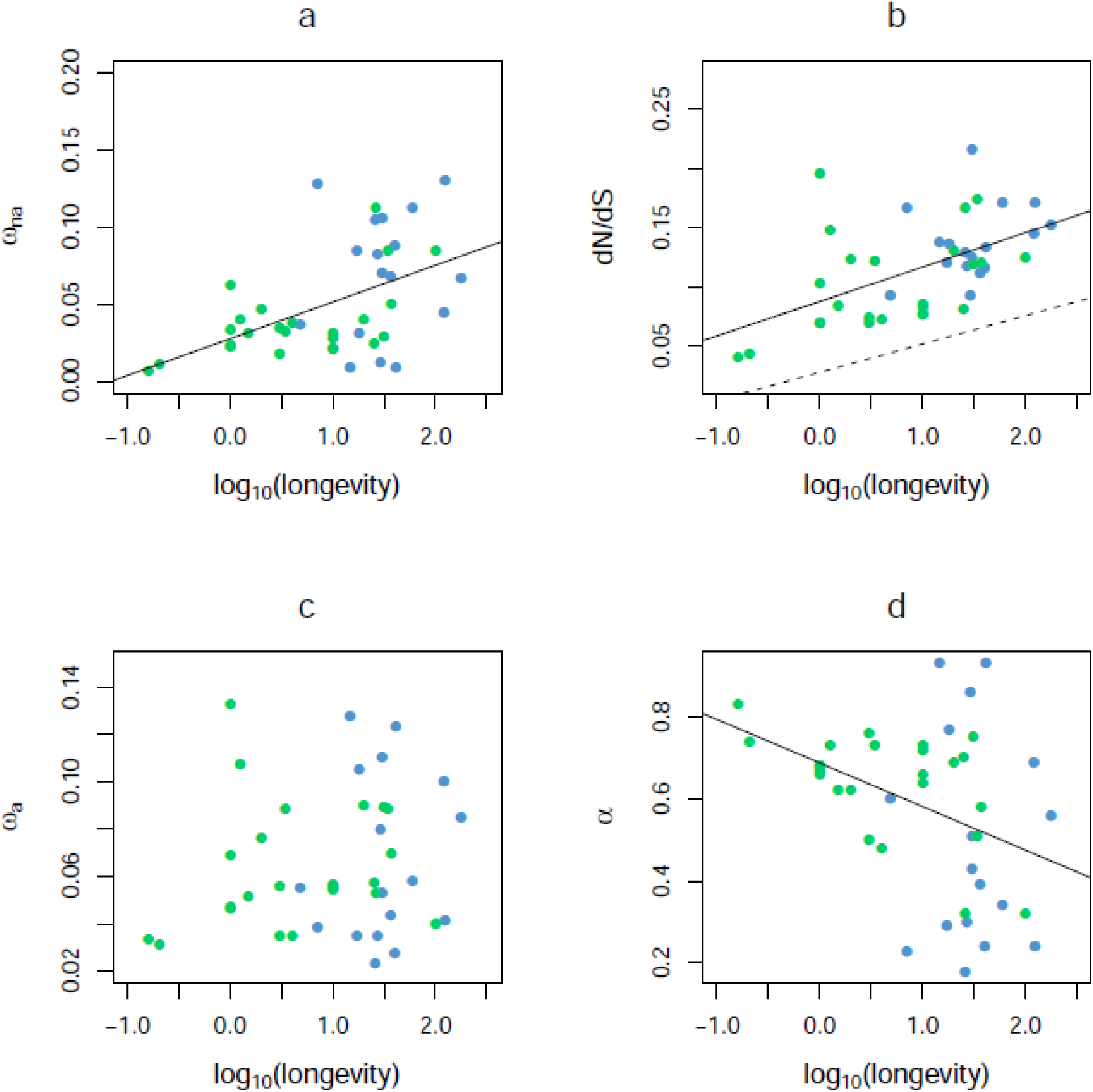
Effect of species longevity on protein divergence parameters. *n*=41 species; blue: vertebrates; green: invertebrates; a: r^2^=0.25, *p*-val<10^−3^; b: r^2^=0.28, *p*-val<10^−3^; c: r^2^=0.02, not significant; d: r^2^=0.15, *p*-val=0.01.

**S3 Figure.**
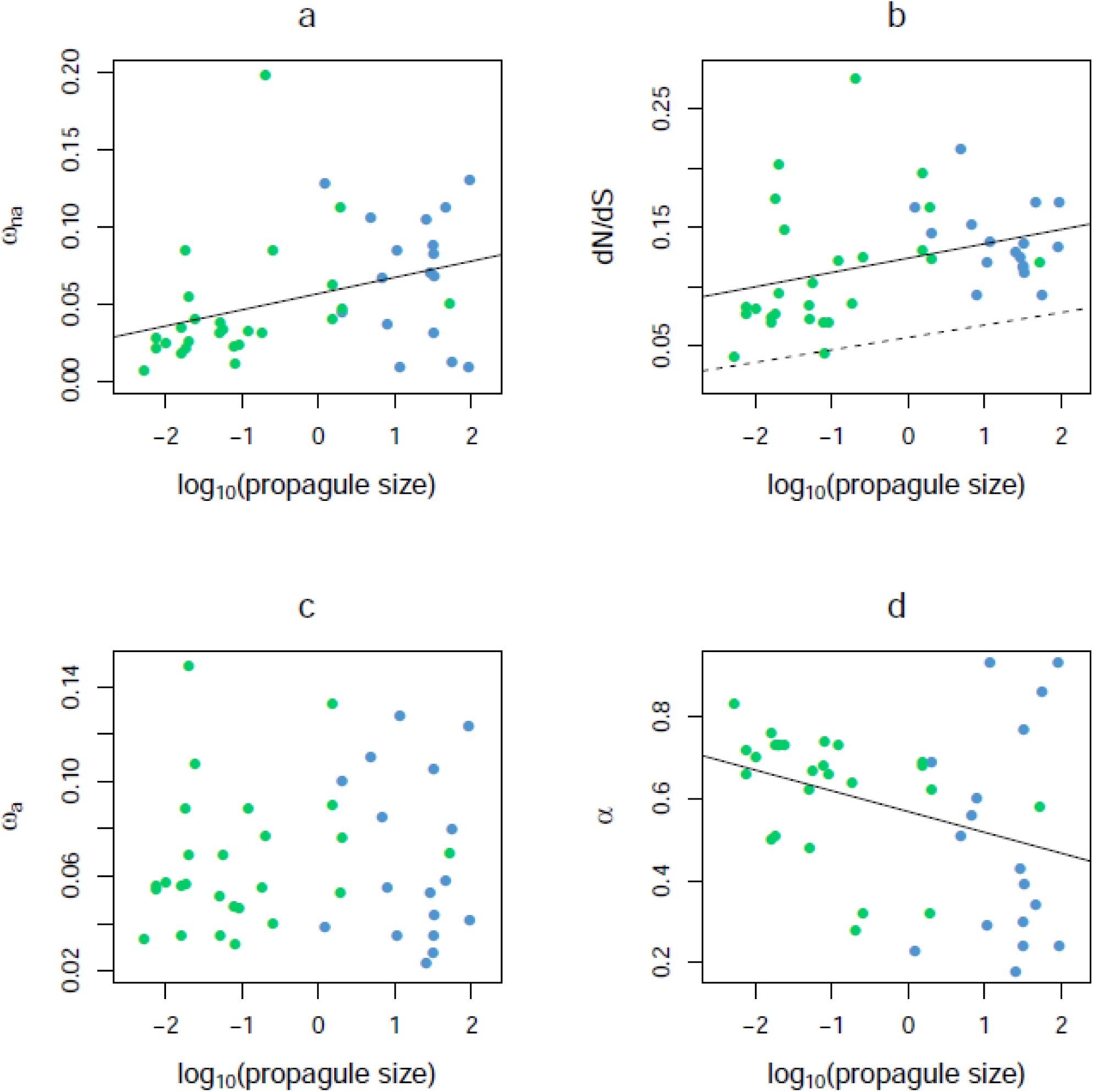
Effect of species propagule size on protein divergence parameters. *n*=43 species; blue: vertebrates; green: invertebrates; a: r^2^=0.13, *p*-val=0.02; b: r^2^=0.12, *p*-val=0.02; c: r^2^=0.005, not significant; d: r^2^=0.12, *p*-val=0.02.

**S4 Figure.**
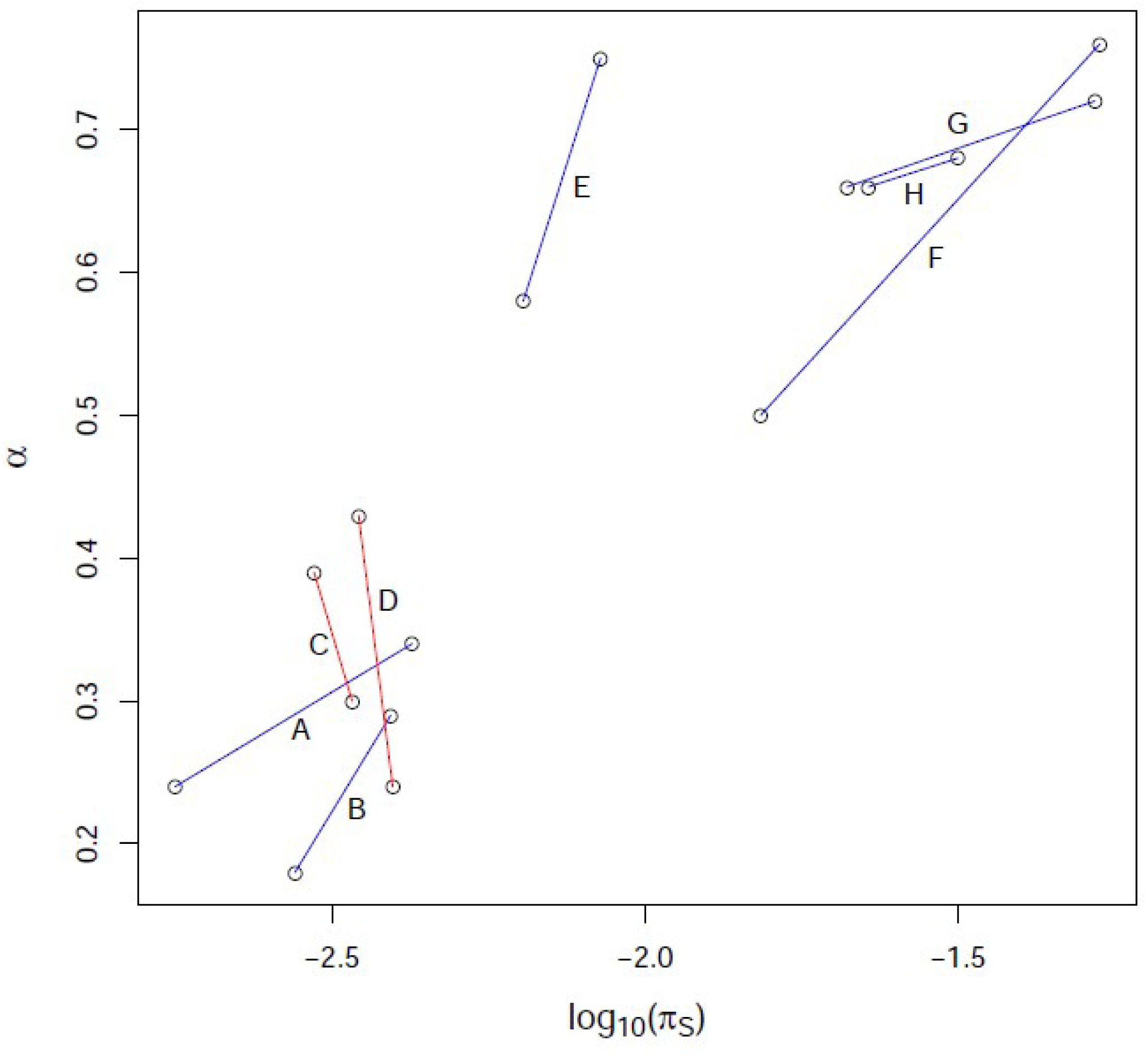
Differences in π_S_ and α between “mirror” species. Segments connect pairs of mirror species. Blue: increasing α with π_S_; red decreasing α with π_S_; A: (*Homo sapiens, Pan troglodytes*); B: (*Nycticebus coucang, Galago senegalensis*); C: (*Eulemur mongoz, Eulemur coronatus*); D: (*Chlorocebus aethiops, Macaca mulatta*); E: (*Varecia variegata variegata*, *Propithecus coquereli*); F: (*Ciona intestinalis* A, *Ciona intestinalis* B); G: (*Echinocardium mediterraneum, Echinocardium cordatum* B2); H: (*Thymelicus sylvestris, Thymelicus lineola*)

**S1 Text. Simulation protocol and results.**

## Supporting Information: S1 Text – Simulation protocol and results

The reliability of the newly introduced methods was assessed by analysing simulated data sets. Forward simulations were performed using SLIM v1.8 [1] in a way very similar to the Messer and Petrov (2013) study [2].

## 1. Constant population size

### 1.1 Simulation protocol

SLIM models mutation, drift, selection and linkage. We simulated the evolution of 250 equally-spaced genes carried by a 10 Mb-long chromosome (one gene every 40 kb). Each gene was made of eight exons of length 150 bp each, separated by introns of length 1.5 kb. Genes were flanked by a 550-bp–long 5’ UTR and a 250-bp–long 3’ UTR. One fourth of the coding and UTR positions were assumed to evolve neutrally, and three fourth were assumed to be under selection. We used a mutation rate of 2.5 × 10^−8^ per site and generation, a recombination rate of 10^−8^, and a panmictic diploid population of constant size *N* = 10^4^. These are identical to the settings of Messer and Petrov (2013).

The selected mutations were codominant and either deleterious or beneficial, the proportion of beneficial ones being called *p*. The selection coefficient of advantageous mutations was fixed to *s_b_*=10^−3^. The selection coefficients of deleterious mutations followed a Gamma distribution of shape 0.3 and mean *s_d_*. *p* and *s_d_* were varied among simulations in order to obtain a wide range of simulated α while keeping the simulated *d*_N/_*d*_S_ similar to that of real data sets.

Simulations were run during 10^6^ generations. Every 10^5^ generations a sample of 7 diploid individuals was taken and allele frequencies at segregating positions were recorded, separately for neutral and selected mutations (simulated SFS). For each allele frequency category, counts were summed across the ten *k*.10^5^th generations. At generation 10^6^, substituted positions were counted, separating neutral, deleterious and beneficial mutations, only counting mutations having appeared after generation 10^5^ (burn-in). This is equivalent to simulating a set of 2,500 genes carried by ten distinct chromosomes during 10^5^ generations. Twelve distinct data sets were generated this way, each using a different (*p*, *s_d_*) pair. Each simulation took ~1.5 day. Three estimates of α and ω_a_ were calculated for each simulated data set, namely the Gamma estimate, the GammaExponential estimate, and the ScaledBeta estimates (see main text). They were compared to the “true” (=simulated) values. These methods assume free recombination between loci and SNPs. The simulation model is therefore different from the inference models.

### 1.2 Results

The number of simulated neutral SNPs was ~3500 per data set, and the number of simulated neutral substitutions ~2800, which is reasonably similar to the real data sets analysed in this study (see S1 Table). The simulated α varied between 0 and 0.63 across data sets, the simulated between 0 and 0.12, and the simulated *d*_N/_*d*_S_ ratio between 0.06 and 0.26. The estimated α and ω_a_ were plotted against the simulated (true) ones:

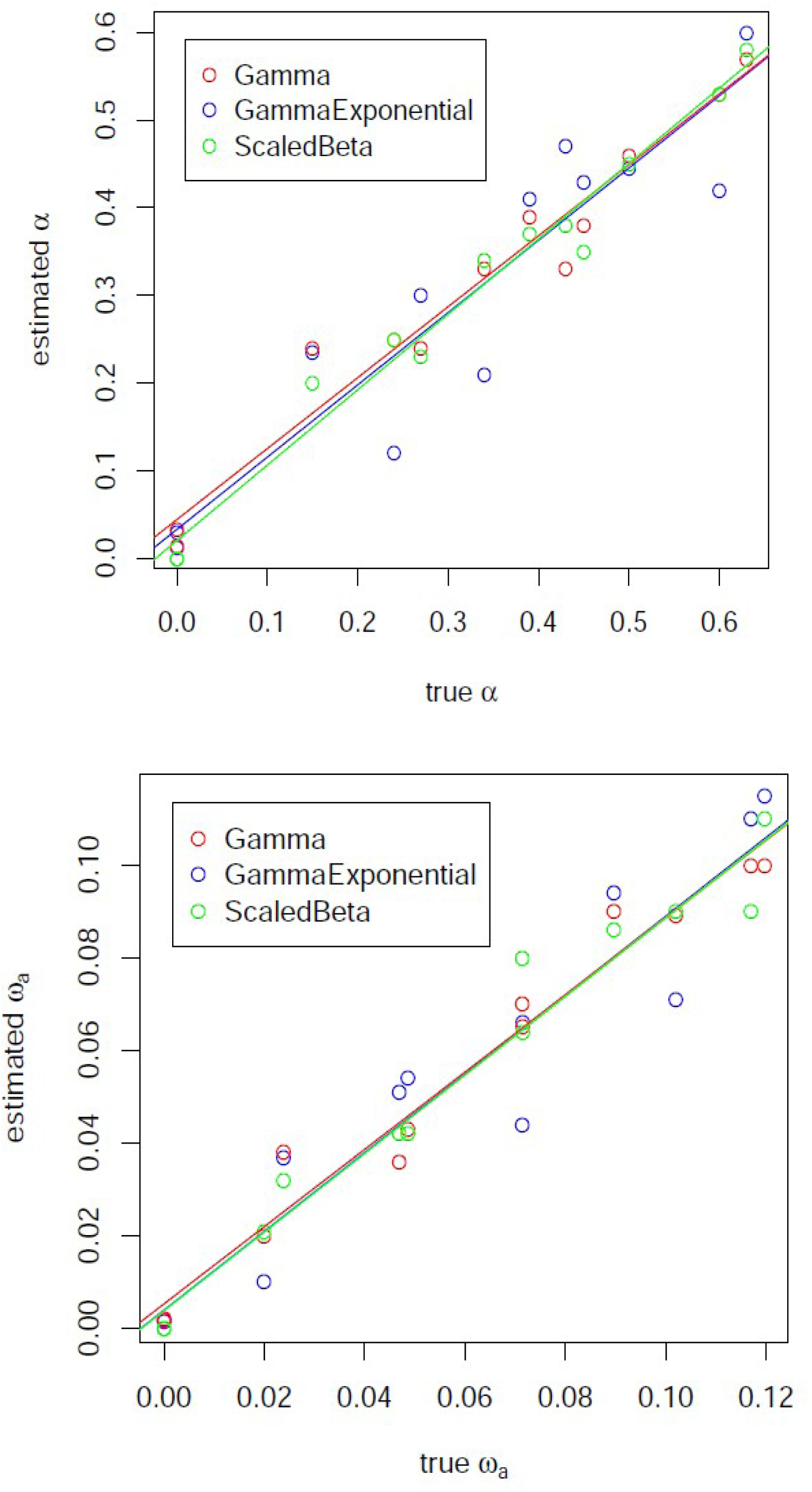

The three estimates performed well; regression lines are almost undistinguishable from the (y=x) line (dotted), and the correlation coefficient between true and estimated values was above 0.9 in all three cases. In all 12 data sets the Gamma model outperformed the GammaExponential and ScaledBeta models according to Akaike’s Information Criterion, which was expected since the simulations assume a Gamma distribution of selection coefficients. The latter two models, although over-parametrized in this specific case, did perform quite well as far as α and ω_a_ estimation were concerned.

These results are in agreement with previous simulation studies, which assessed the accuracy of distinct but related estimators of α. Performing simulations very similar to ours, Messer and Petrov (2013) [2] showed that the Keightley & Eyre-Walker (2007) method [3] returns reasonably good estimates of α in spite of the distortion of SFS’s due to linked selection – even though demographic parameters can take irrelevant values in such cases. Eyre-Walker and Keightley (2009) [4] compared the Eyre-Walker et al. (2006) [5] and the Keightley & Eyre-Walker (2007) [3] methods based on real data and simulations. They found that the former, which is very similar to the Gamma estimate of this study, performed a bit better than the latter, both being quite robust to departure from the assumption of independence between SNPs. The simulation scheme of Eyre-Walker and Keightley (2009) [4] did not incorporate any adaptive effect.

## 2. Fluctuating population size

A second round of simulations was conducted assuming varying effective population (*N*) in time. *N*(*t*) was assumed to follow a Markov process, the average waiting time between two events of population size change being 50,000 generations. After each event of *N* change, the new *N* was randomly drawn in the [5,000; 50,000] interval, in such a way that 1/*N* was uniformly distributed over [1/50,000; 1/5,000]. The average *N* was close to 10^4^.

The deleterious effect *s_d_* was set to −0.25, and three positive selection regimes were simulated: *p*=0 (no adaptive mutation), *p*=0.0015 (medium), *p*=0.007 (high adaptive rate). Other parameters were identical to section 1 above. Again, despite some additional sampling variance, the true and estimated α and ω_a_ were well correlated, suggesting that DFE-based McDonald-Kreitman methods are reasonably robust to fluctuations in effective population size (see figures below).

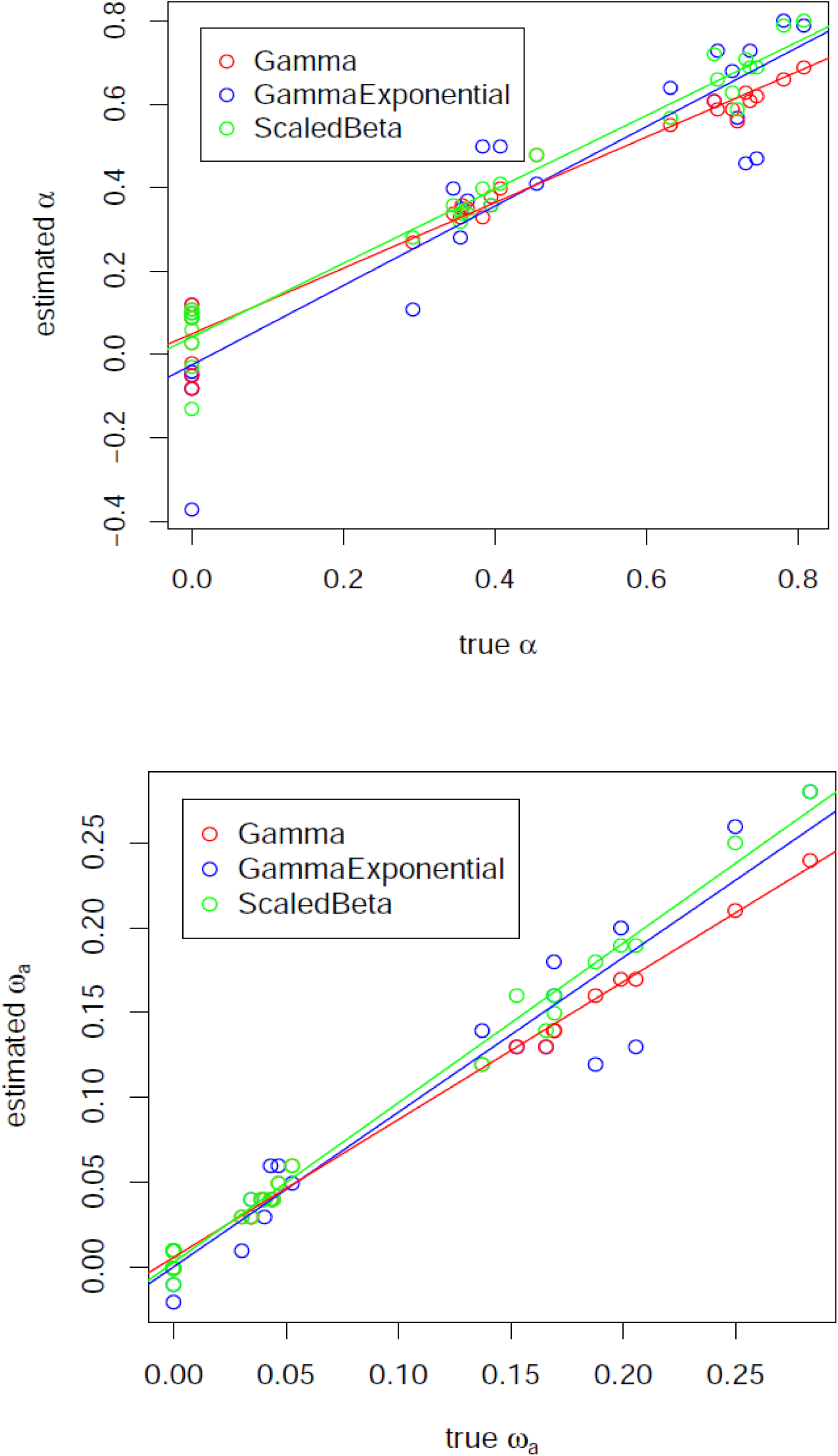

